# Bacterial association with metals enables *in vivo* tracking of microbiota using magnetic resonance imaging

**DOI:** 10.1101/2022.05.05.490857

**Authors:** Sarah C. Donnelly, Neil Gelman, R. Terry Thompson, Frank S. Prato, Jeremy P. Burton, Donna E. Goldhawk

## Abstract

Bacteria constitute a significant part of the biomass of the human microbiota, but their interactions are complex and difficult to replicate outside the host. Exploiting the superior resolution of magnetic resonance imaging (MRI) to examine signal parameters of selected human isolates may allow tracking of their dispersion throughout the body. We investigated longitudinal and transverse MRI relaxation rates and found significant differences between several bacterial strains. Common commensal strains of lactobacilli display notably high MRI relaxation rates, partially explained by outstanding cellular manganese content, while other species contain more iron than manganese. *Lactobacillus crispatus* show particularly high values, 4-fold greater than any other species; over 10-fold greater signal than relevant tissue background; and a linear relationship between relaxation rate and fraction of live cells. Different bacterial strains have detectable, repeatable MRI relaxation rates that in future may enable tracking of their persistence in the human body for enhanced molecular imaging.

IMPORTANCE

To understand how spatial and temporal distribution of microbiota impact human health, dynamic tools for monitoring microbiota landscapes inside the host are needed. Particularly when considering the complexity of the gastrointestinal tract and the microbiota that dwell within, tools for monitoring deep segments of the gut non-invasively are required. Medical imaging provides solutions that enable the study of microorganisms in their preferred niche regardless of health status. To bootstrap this technology, we investigated the magnetic resonance imaging (MRI) properties of bacterial isolates and showed that outstanding signal detection is an inherent property of several strains. Among these, we showed that bacteria relying on manganese metabolism have an MRI characteristic that is distinct from mammalian cells. Our findings will lead to direct and safe imaging of bacteria; influence how we monitor both infection and gut health; and help direct the use of antibiotics to curtail the growing threat of antibiotic resistance.

## Introduction

Microbial-host interactions are widespread and many of these are complex symbiotic relationships as seen in the human microbiota at different sites in the body, though most occur in the oral cavity, and intestinal and reproductive tracts ^1^. These interactions cause changes in metabolism and signaling pathways, not only in health but also in disease. The microbiota serves many functions, including protection from pathogens, production of vitamins and metabolites for energy, and major roles in homeostasis ^2, 3^. An understanding of the importance of the microbiota in many diseases has been rapidly increasing, with ramifications for autoimmune diseases (*e.g.* rheumatoid arthritis ^4^, type 1 diabetes ^5^); metabolic syndromes (*e.g.* type 2 diabetes ^6, 7^, coronary artery disease ^8^); and neuropsychiatric disorders (*e.g.* depression ^9^, psychosis ^10^).

Microbiota interactions are complex and difficult to replicate outside of the host. Despite this, the current gold standard for study of the composition, properties and functions of microbiota entails *ex vivo* analyses, involving removal of microbial samples from the mucosa and propagation *in vitro* for high-throughput techniques like 16S rRNA gene sequencing ^11^, shotgun metagenomics ^12^, or metabolomics ^13^. These approaches are susceptible to sample degradation, contamination, or potentially undergoing change(s) in bacterial growth that subsequently misrepresent the microbiota of the original sample. The microbiota within feces, for example, is not representative of the intestinal mucosa ^14–16^. Since the survival of many microbes depends on bacterial-host interactions, specific nutrient requirements and atmospheric conditions, faithfully replicating the *in vivo*environment in *ex vivo* cultures is difficult.

Medical imaging platforms offer the possibility of studying biological events *in vivo*, including the microbiota. However, the ability to label microorganisms other than for research purposes has been challenging ^17^. Some new labels have been developed that are specifically utilised by certain microbes, for example 2-deoxy-2-[^18^F]fluoro-D-sorbitol for the detection of *Enterobacterales* using positron emission tomography (PET) ^18^. Given this potential of nuclear medicine for streamlining medical treatment, *ex vivo* methods to assess microbiota might best be used as tools to validate *in vivo* molecular imaging.

Despite widespread clinical use of magnetic resonance imaging (MRI), very little research has employed this technology to specifically detect bacteria ^19–23^, and of these reports, virtually all relied on cell labels for contrast enhancement. To date, most MRI studies image bacterial infection indirectly by focusing on host inflammation ^24, 25^. Even if bacteria could be directly detected at clinically relevant levels, questions remain about whether the bacterial magnetic resonance (MR) signal can be resolved from surrounding mammalian tissue based on differences in relaxation rates, diffusion, or other cellular MR characteristics. Relevant to this, the association of various bacterial types with metals both in the environment and in medicine has long been appreciated ^26–28^. Ferromagnetic and paramagnetic metals like iron, manganese, nickel, cobalt, gadolinium, and dysprosium influence MRI ^29^. Indeed, bacterial acquisition of metal cofactors like iron and manganese is widely considered a form of bacterial pathogenesis, with many siderophores encoded on pathogenicity islands to improve survival in low iron environments ^30, 31^. In gram-negative uropathogenic *Escherichia coli* (UPEC), iron and manganese uptake are linked by expression of the *sitABCD* operon within their pathogenicity island(s) ^32^. The genes in this operon include those encoding iron and manganese uptake proteins, regulated by transcription factors involved in the ferric uptake regulator (Fur) system as well as the manganese transport regulator (MntR) ^33^. Unlike many bacteria, lactobacilli do not use iron as a cofactor, relying instead on manganese ^34, 35^ and its active import through the manganese/cadmium transporter MntA ^36, 37^. Thus, bacterial acquisition and storage of ferromagnetic iron or paramagnetic manganese are important factors to consider in the context of imaging bacteria by MRI.

Recognizing the importance of microbiota in human health; the need for *in vivo* visualization of microbial factors that underlie disease manifestation and progression; and the superior resolution of MRI for examining soft tissues ^38^, we sought to provide evidence on whether bacteria may be detected in the future with MRI, either alone and/or in combination with hybrid modalities like PET/MRI. Given large genetic variability and differences in iron and manganese handling ^39^, we hypothesized that MR measures could be quite different than typical tissue values and could vary drastically between bacterial species and strains. To explore these differences, we examined MR relaxation rates of various pathogenic, probiotic, and commensal bacteria. Here we show that MR relaxation rates vary among bacteria partially due to differences in metal handling. In particular, lactobacilli have high MR relaxation rates which correlate to fraction of cells in the MR volume. These characteristics make certain species ideal candidates for developing methods to track bacteria *in vivo* using MRI.

## Results

### MR relaxation rates of bacteria

In healthy females, the lower urinary system is dominated by species of lactobacilli ^40^ and mostly by strains of *Lactobacillus crispatus* ^41^. Our study of species detected in the female urogenital system, which typically has relatively low microbial diversity, may offer a more simplistic microbiota for *in vivo* imaging. Other microbes of interest were included for MRI signal comparisons to probiotic, pathobiont, pathogenic and commensal strains. *Staphylococcus aureus* have been characterized for their acquisition and handling of iron, a ferromagnetic MRI detectable metal ^27^. *Escherichia coli* are interesting for MRI study owing to significant genetic polymorphisms at the strain level, bestowing characteristics representative of their adaptation to diverse habitats and their roles as commensals and pathogens, including UPEC.

Bacteria were cultured and analyzed in a MRI cell phantom (Fig. 1, a, b and c). In this subset of bacteria, MR relaxation rates were variable (Fig. 1, d, e and f) but frequently higher than recorded in mammalian cell types ^42, 43^. Transverse relaxation rates of bacteria were dominated by the R2 component (paired t-test, t=1.395 df=31 P=0.1729) and these measures in *Lactobacillus gasseri* ATCC33323 were significantly higher than other species examined (Fig. 1, d and e). Both *Lactobacillus rhamnosus* GR-1 and *E. coli* BL21(DE3) displayed higher R2* transverse relaxation rate than *Proteus mirabilis* 296 and *Staphylococcus aureus* Newman (Fig. 1d, ANOVA F(11,28)=19.82). Remarkably, while multiple *Lactobacillus* spp. demonstrated high transverse relaxation rates (1/T2, Fig. 1e, ANOVA F(11,28)=23.77), the signal intensity of *L. crispatus* ATCC33820 decayed to background before the shortest echo time (TE, 13 ms), thus preventing an accurate measurement in undiluted samples. This finding confirmed that select microbes may be amenable to cell tracking by MRI owing to relaxation rates that far exceed those in mammalian tissue.

**Figure 1.**
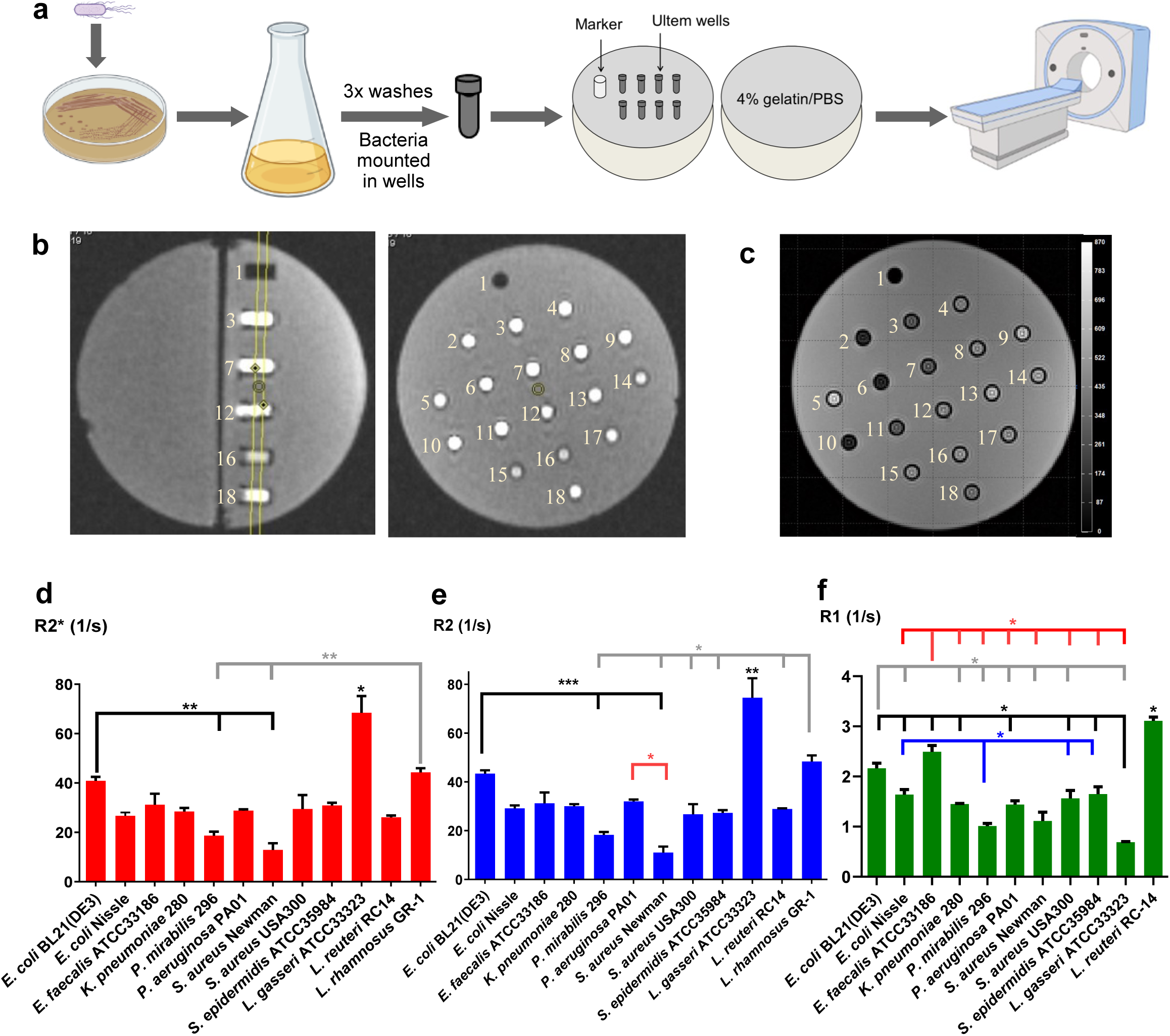
MRI relaxation rates of different bacterial species vary widely. **a)** Cultures from different bacterial species were harvested, washed, and loaded into Ultem wells prior to mounting in one hemisphere of a gelatin phantom. Hemispheres were then assembled to form a 9 cm spherical cell phantom; placed in a knee coil; and scanned at 3T by MRI. **b)** Locator images show both sagittal (left) and axial (right) cross-sections through a gelatin cell phantom. The MR signals for relaxation measurements were acquired from a 3 mm slice (yellow lines, left panel) passing through all wells. Numbers to the left of each well designate sample identity. In this example, 1 is a plastic marker; wells 2-13 contain dilutions of *L. crispatus* ATCC33820 in gelatin, where 2, 6 and 10 are 1/2 dilutions; 3, 7 and 11 are 1/4 dilutions; 4, 8 and 12 are 1/8 dilutions; and 5, 9 and 13 are 1/16 dilutions; wells 14 – 18 contain *S. aureus* and *E. coli* mutants. **c)** In a representative T2-weighted spin echo image, signal intensity is displayed at echo time (TE) 20 ms, with corresponding scale bar in arbitrary units. **d-f)** In all cases, the total transverse relaxation rate R2* (d, red bars) is dominated by the R2 component of transverse relaxation (e, blue bars). *Lactobacillus gasseri* displayed significantly higher R2* and R2 than any other species tested. *Escherichia coli* BL21(DE3) and *L. rhamnosus* displayed higher transverse relaxation than several species (d and e, black and gray lines, respectively). *Pseudomonas aeruginosa* displayed higher R2 than *S. aureus* Newman (e, red line). Longitudinal relaxation rates also varied significantly between bacterial species (f, green bars), with *L. reuteri* displaying higher R1 than all other species. R1 of *E. coli* BL21(DE3) and *E. faecalis* were higher than most other species (f, gray and red lines, respectively). *Lactobacillus gasseri* and *P. mirabilis* both had lower R1 than several other bacterial species (f, black and blue lines, respectively). Bar graphs show the mean +/- SEM (n = 3–5). *, p < 0.05; **, p < 0.01; ***, p < 0.001.

Longitudinal relaxation rates (R1) reflecting spin-lattice interactions were also distinct, with lactobacilli paradoxically among the highest and lowest recorded in this study (Fig. 1f, ANOVA F(10,24)=41.84). For *Lactobacillus reuteri* RC-14, R1 (3.11 ± 0.08 s^-1^) was significantly higher than all other species examined while for *L. gasseri* ATCC33323, R1 (0.69 ± 0.01 s^-1^) was lower than most. These data suggest that R1 in addition to R2* and R2 measurements may be useful for differentiating some lactobacilli from other bacterial species.

To position measures of bacterial relaxation rates in clinical context, *in vivo* MRI of the human bladder was acquired to establish appropriate tissue background for urinary strains. Both T1 and T2* maps from male (Fig. 2, a and b) and female (Fig. 2, c and d) volunteers provided similar values for R1 (1/T1) and R2* (1/T2*; Supplementary Table 1) in regions of interest contouring the bladder wall (Fig. 2 arrows). A plot of R1 vs R2* comparing the five groups of bacteria indicates the relationship between family members and MR measures (Fig. 2e). In most species examined, either R1 or R2* relaxation rates are well above tissue background, as assessed in the healthy human bladder. As expected, the same is true for R1 vs R2 (Supplementary Fig. 1).

**Figure 2.**
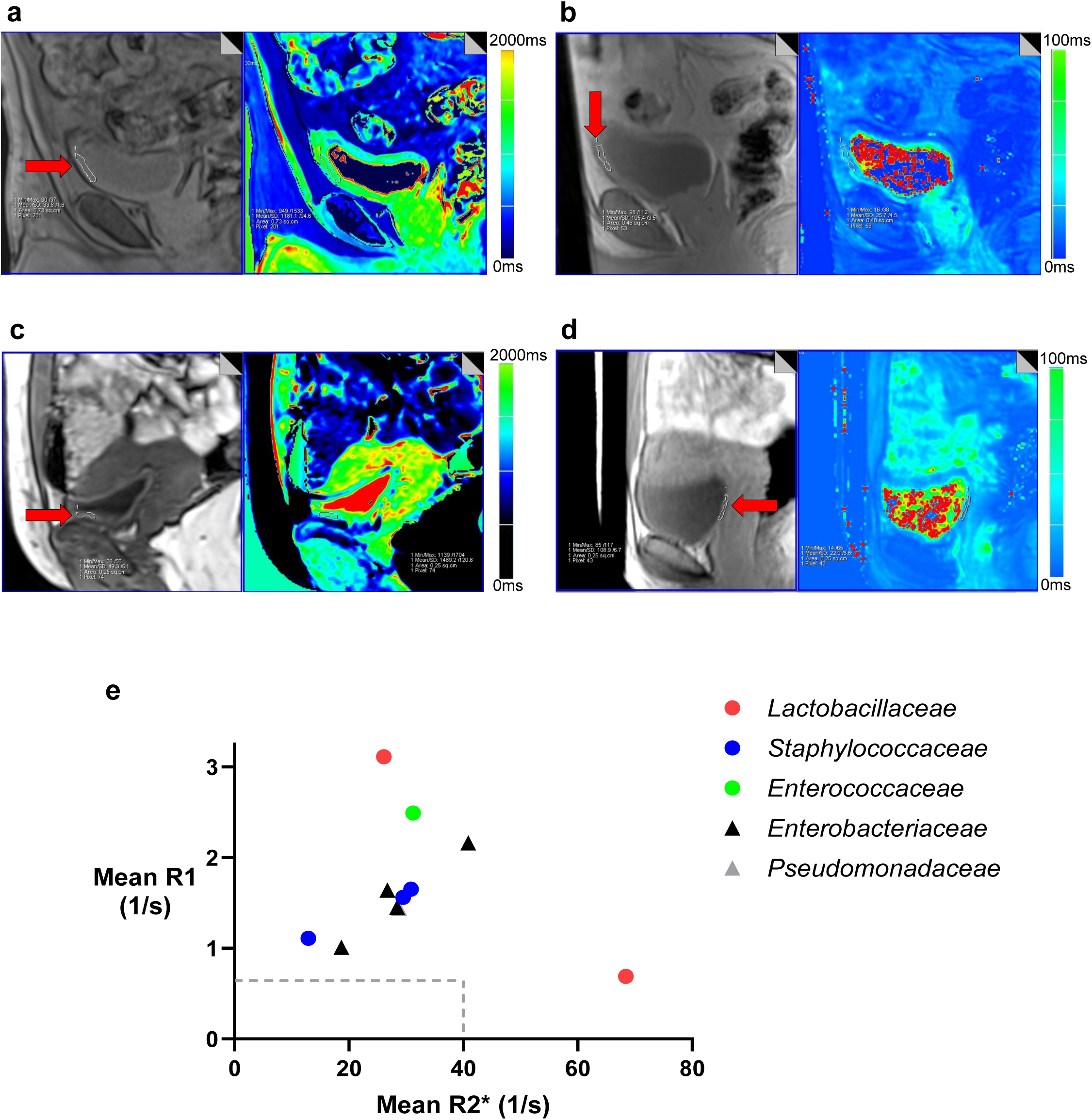
Relaxation rates of urinary bacterial isolates exceed *in vivo* measures of the healthy human bladder. **a-d)** Representative MR images of the healthy bladder were acquired at 3T in male (a and b) and female (c and d) volunteers (n = 3). T1 images and maps (a and c) and T2* images and maps (b and d) show a sagittal section of the bladder with region of interest along the bladder wall outlined in white, as shown by the red arrows. Corresponding heat maps (right panels) indicate the relaxation times with matching scale bars. **e)** A scatter plot of R1 vs R2* shows the mean relaxation rate measurements of bacteria (n = 3–5). Gram positive species are denoted by circles and gram-negative species by triangles. Bacteria are grouped into their respective families as indicated by symbol colour: red, *Lactobacillaceae*; blue, *Staphylococcaceae*; green, *Enterococcaceae*; black, *Enterobacteriaceae*; gray, *Pseudomonadaceae*. Broken gray lines show average R1 (0.67 s^-1^) and R2* (39.3 s^-1^) in the healthy human bladder.

### Fe and Mn quantification in various bacterial species

To better understand diverse MRI measures in urinary isolates, the influence of metal co- factors was examined. After cell lysis and quantification of total cellular protein (Fig. 3a), different bacterial strains were analyzed by inductively-coupled plasma mass spectrometry (ICP-MS) to quantify the total cellular content of iron (Fig. 3b) and manganese (Fig. 3c). Iron was below detectable levels in all samples of *Lactobacillus* spp. examined (Fig. 3b), consistent with their preference for manganese ^34^. Thus, by comparison, most bacterial species contained significantly more iron than lactobacilli (black line, p < 0.05; Kruskall-Wallis P < 0.001). *Staphylococcus aureus* Newman contained more iron than *E. coli* Nissle and *Enterococcus faecalis* ATCC33186 (gray line). In addition, *Klebsiella pneumoniae* 280 contained more iron than *E. faecalis* (red line).

**Figure 3.**
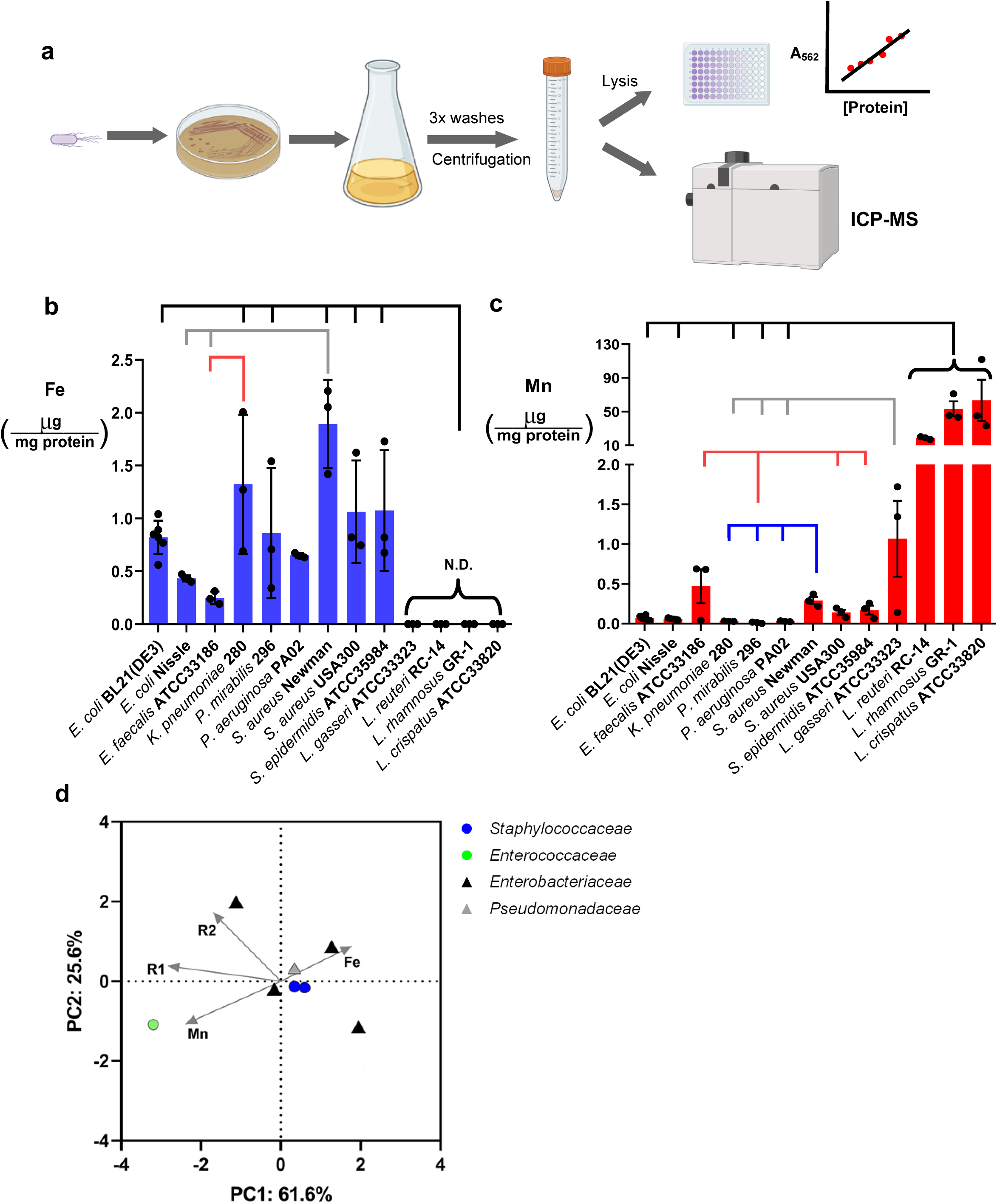
Total cellular iron and manganese vary widely between bacterial species. **a)** Cultured bacteria were washed and pelleted prior to lysis and protein quantification followed by ICP-MS to measure elemental iron and manganese. **b-c)** Total cellular iron and manganese content was normalized to total cellular protein. Both iron (b, blue) and manganese (c, red) content varied between bacterial species. Iron content of most bacterial species was significantly higher than that of all lactobacilli examined (b, black line) while *S. aureus* Newman contained more iron than *E. coli* Nissle and *E. faecalis* (b, gray line). In addition, *K. pneumoniae* contained more iron than *E. faecalis* (red line). In panel c, *P. mirabilis* contained less Mn than *E. faecalis* (red line) and all staphylococci examined (red and blue lines). *Staphylococcus aureus* Newman also contained more Mn than *K. pneumoniae* and *P. aeruginosa* (c, blue line); whereas all Lactobacillus samples contained more Mn than many other bacterial species examined (c, black and gray lines). Data are displayed as individual values (black circles) with mean +/- SEM (n = 3-6) and all comparisons reflect p < 0.05. (N.D., not detectable). **d)** Principal component analysis of mean MR and ICP-MS measures in bacteria, excluding outliers. The distance between samples on the plot represents differences in bacterial metal handling and MR measures, with 87.2% of total variance being explained by the first two components shown. The association of variables are depicted by the direction of the gray arrows. Each coloured point represents a separate bacterial strain or species as the mean of 3-5 replicates of each variable measurement. Points are coloured by bacterial family: blue *Staphylococcaceae*; green, *Enterococcaceae*; black, *Enterobacteriaceae*; gray, *Pseudomonadaceae*. Gram positive species are denoted by circles and gram-negative by triangles.

Total cellular content of elemental manganese varied considerably between bacteria, ranging between 0.0034 – 112 µg Mn / mg protein (Fig. 3c). *Staphylococcus aureus* Newman contained more manganese than *K. pneumoniae*, *P. mirabilis*, and *Pseudomonas aeruginosa* (blue line; Kruskall-Wallis P < 0.001). *Proteus mirabilis* 296 contained the lowest manganese content (red line) while all other significant differences were due to the high manganese content of lactobacilli. *Lactobacillus reuteri* RC-14, *L. rhamnosus* GR-1, and *L. crispatus* ATCC33820 had higher manganese than all non-*Lactobacillus* species except for staphylococci and *E. faecalis* (black line). *Lactobacillus gasseri* ATCC33323 had significantly more manganese than *P. aeruginosa, K. pneumoniae,* and *P. mirabilis* (gray line).

These data show that both iron and manganese levels vary widely between different bacterial species, raising the possibility that manganese rather than iron may contribute extensively to the high MR relaxation rates of *Lactobacillus* spp. Note however that *L. gasseri* ATCC33323, with over 10-fold less manganese than the other lactobacilli examined, exhibited a mean R2 of 74.53 ± 8.00 s^-1^, second only to *L. crispatus* ATCC33820 (discussed below). Hence elemental manganese content is not the only determinant of R2.

In a principal component analysis evaluating mean R2, R1, elemental iron and manganese content, two principal components account for 87.2% of the variance. No significant correlations between variables are apparent but species of lactobacilli appear as outliers (Supplementary Fig. 2 and Supplementary Table 2) based on extraordinary R2 in the case of *L. gasseri* and both R1 and manganese content in the case of *L. reuteri*. *Staphylococcus aureus* Newman also represents an outlier owing to elevated iron content and low R2. Removing these outliers to reduce bias in the principal component analysis, confirms the positive correlations between bacterial R1 and manganese content (Fig. 3d, Supplementary Table 3).

### MR relaxivity of *L. crispatus* ATCC33820

To examine the outstanding MR signal intensity of *L. crispatus* ATCC33820, given its rapid T2 decay, samples were serially diluted in 4% gelatin / PBS prior to mounting in the gelatin cell phantom (Fig. 4a). Following a 1/2 dilution, signal intensity still decayed to background levels by the third TE (25 ms) but R2 and R2* were nevertheless measurable (Fig. 4b; ANOVA for R2*: F(4,30)=35.69 and for R2: F(4,30)=53.47). Indeed, transverse relaxation rates were more reliable as the dilution factor increased and more TE could be used for decay curve fitting. With this signal advantage, *L. crispatus* may be detected by MRI even when relatively few cells are present. To explore this, colony forming units (CFUs) were quantified in *L. crispatus* samples at each dilution, with approximately 4 billion live cells in a volume of 38 mm^3^ (1/32 dilutions) providing measurable signal. At 1/8 dilutions, with approximately 11 billion live cells, R2* and R2 were 71.3 ± 6.10 s^-1^ and 56.6 ± 3.50 s^-1^, respectively, well above typical measures for mammalian cells ^42, 43^ and average measures in relevant tissue like the human bladder (Fig. 2). Moreover, R1 values (Table 1) obtained from 11 billion CFUs or more were also significantly higher than samples with fewer live cells (1/8 vs 1/16 dilutions, p < 0.001; ANOVA F(3,21)=44.08), pointing again to a threshold above which these bacteria might be distinguished from tissue background.

**Figure 4.**
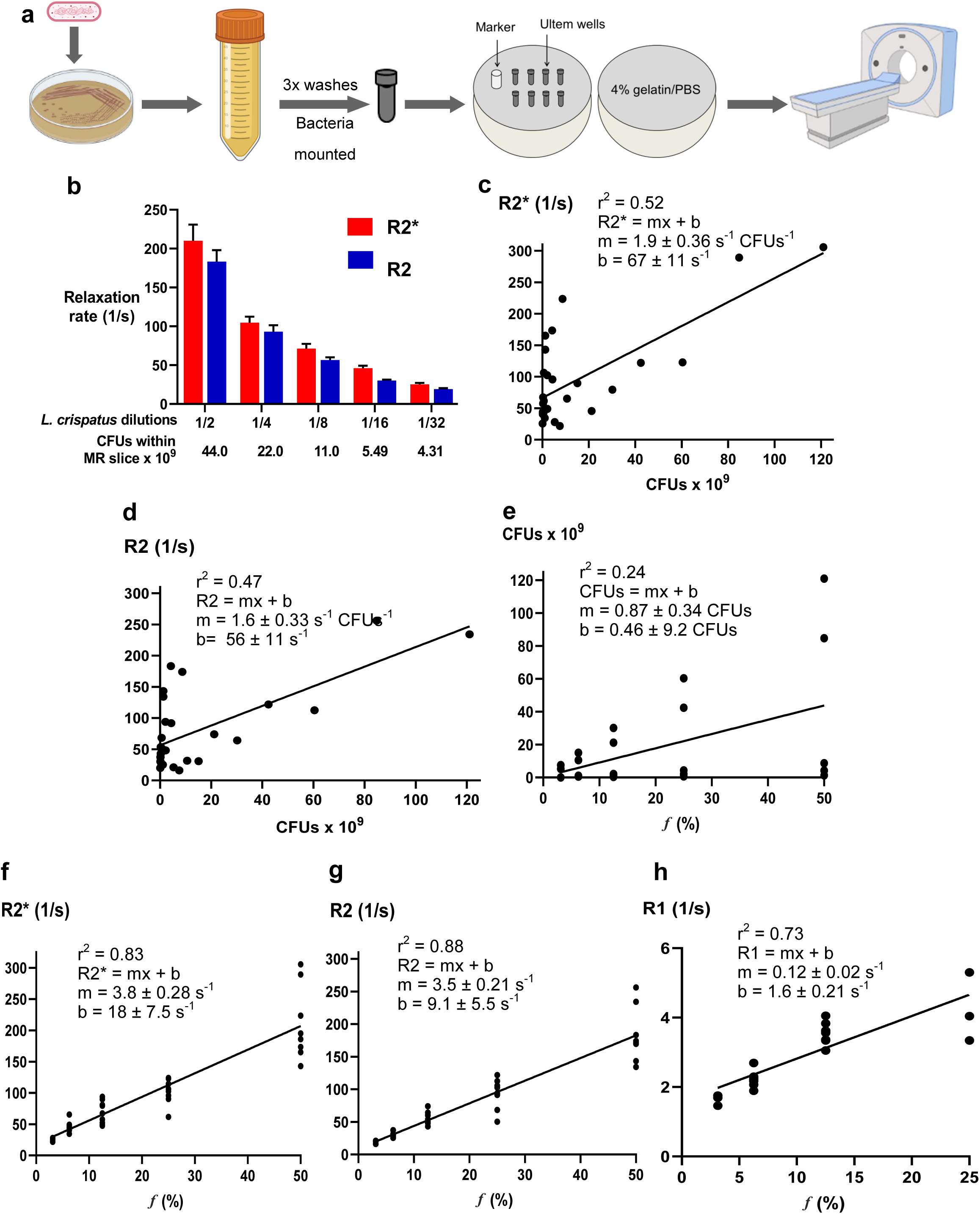
*Lactobacillus crispatus* ATCC33820 display high MR relaxation rates related to amount of live cells. **a)** *Lactobacillus crispatus* were cultured anaerobically, and cells were serially diluted in gelatin/PBS prior to loading into Ultem wells mounted in a gelatin phantom for MRI at 3T. **b)** *Lactobacillus crispatus* displays high transverse relaxation rates when diluted in gelatin / PBS. Bar graphs show the mean +/- SEM (n = 4-10) of R2* (red) and R2 (blue). Colony forming units (CFUs) present within the MR slice were calculated based on CFUs estimated from initial undiluted bacterial cultures **c-d)** Plots of R2* (c) and R2 (d) show individual MR measures as a function of number of live cells (expressed as CFUs) within each MR slice. For both R2* and R2, Pearson’s line of best fit provides a moderately positive linear correlation between CFUs and transverse relaxation rates (p < 0.001). **e)** A scatter plot shows CFUs within the MR slice of individual samples (21 voxels in a 3 mm slice through the well is approximately 38 mm^3^) as a function of the percentage of cells (*f*) loaded into the wells after serial dilution in gelatin/PBS. The line represents a moderate, positive linear correlation between CFUs and fraction of cells (p < 0.05). Irrespective of dilution factor, there is a range in the estimated number of live cells that may contribute to the MR signal in any given well of the cell phantom, with average CFUs indicated by the line of best fit. **f-h)** Fraction of *L. crispatus* cells (*f*) is strongly correlated to R2* (f), R2 (g) and R1 (h) measurements (p < 0.001).

**Table 1.**
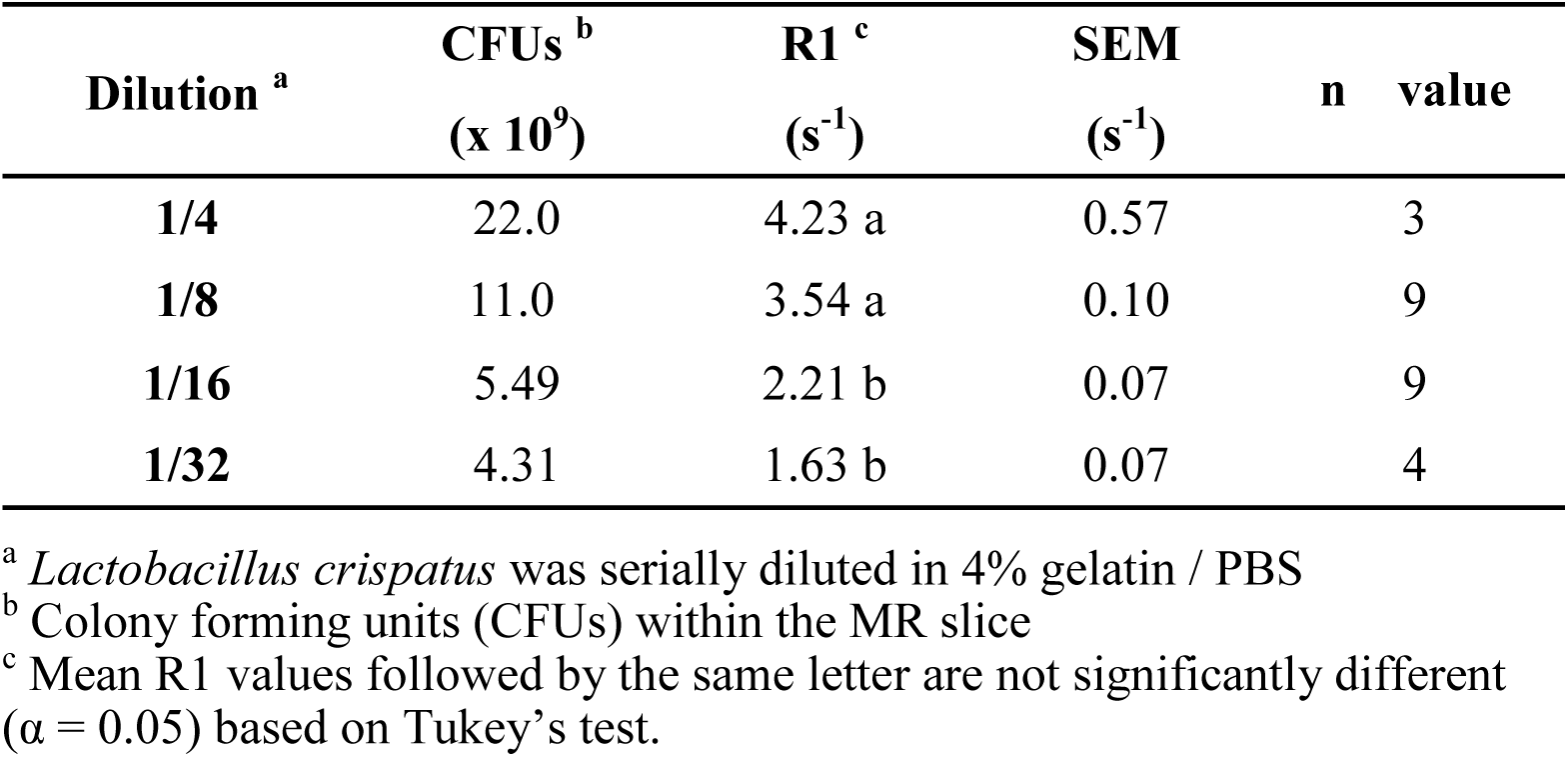
R1 relaxation rates for serially diluted L. crispatus

Since number of CFUs per unit volume varies with each replicate, we explored the correlation between CFUs in the MR slice (where the signal is acquired) and its relaxation rate. Both R2* and R2 show moderate positive linear correlations to the number of live cells (Fig. 4, c and d, respectively; p < 0.001). Thus, as the number of live cells in the region of interest (ROI) increases, so do the MR measures. Based on the moderate, positive linear correlation between CFUs in the slice and fraction of cells, we can estimate the average number of CFUs required to fill the MR volume at any dilution factor (Fig. 4e). However, unlike the variable nature of estimating CFUs, R2*, R2 and R1 of *L. crispatus* ATCC33820 were strongly correlated to the fraction (*f*) of cells in the sample (Fig. 4, f, g and h, respectively). Based on these strong positive correlations, we extrapolated the linear regression equations to estimate that R2* for undiluted *L. crispatus* (*f* = 100 %) is approximately 398 ± 36 s^-1^, R2 is approximately 359 ± 27 s^-1^, and R1 is approximately 13.6 ± 2.2 s^-1^. These estimated values are at least 4-fold higher than any other species examined. Overall, these *L. crispatus* dilution experiments demonstrate that a commensal bacterium of the urogenital tract may be a good target for developing *in vivo* imaging of bacteria using MRI.

### MRI in a population of bacterial and mammalian cells

To explore the potential for distinguishing MR relaxation rates in two distinct cell types within a single MR slice, we examined homogeneous mixtures of human 5637 bladder cells and *L. crispatus* ATCC33820 (Fig. 5a): two cell types that line the urinary tract, have documented interaction ^44^, and whose proximity may influence voxel by voxel analysis of MRI signals. Decay curves of samples at varying ratios show that transverse relaxation rates in these cell mixtures are mono-exponential (Supplementary Fig. 3), indicating that these bacterial and mammalian MR signals cannot be resolved using the standard MR sequences applied here. In addition, R2* values of these mammalian and bacterial cell mixtures are similar to those of *L. crispatus* alone, diluted in gelatin. However, R2 values are markedly lower than those measured in bacterial cell/gelatin phantoms (Fig. 4b vs 5b). While *L. crispatus* diluted in gelatin has a very small R2′ component (R2* − R2, Fig. 4b), in the presence of bladder cells R2′ contributes to approximately half of the R2* value.

**Figure 5.**
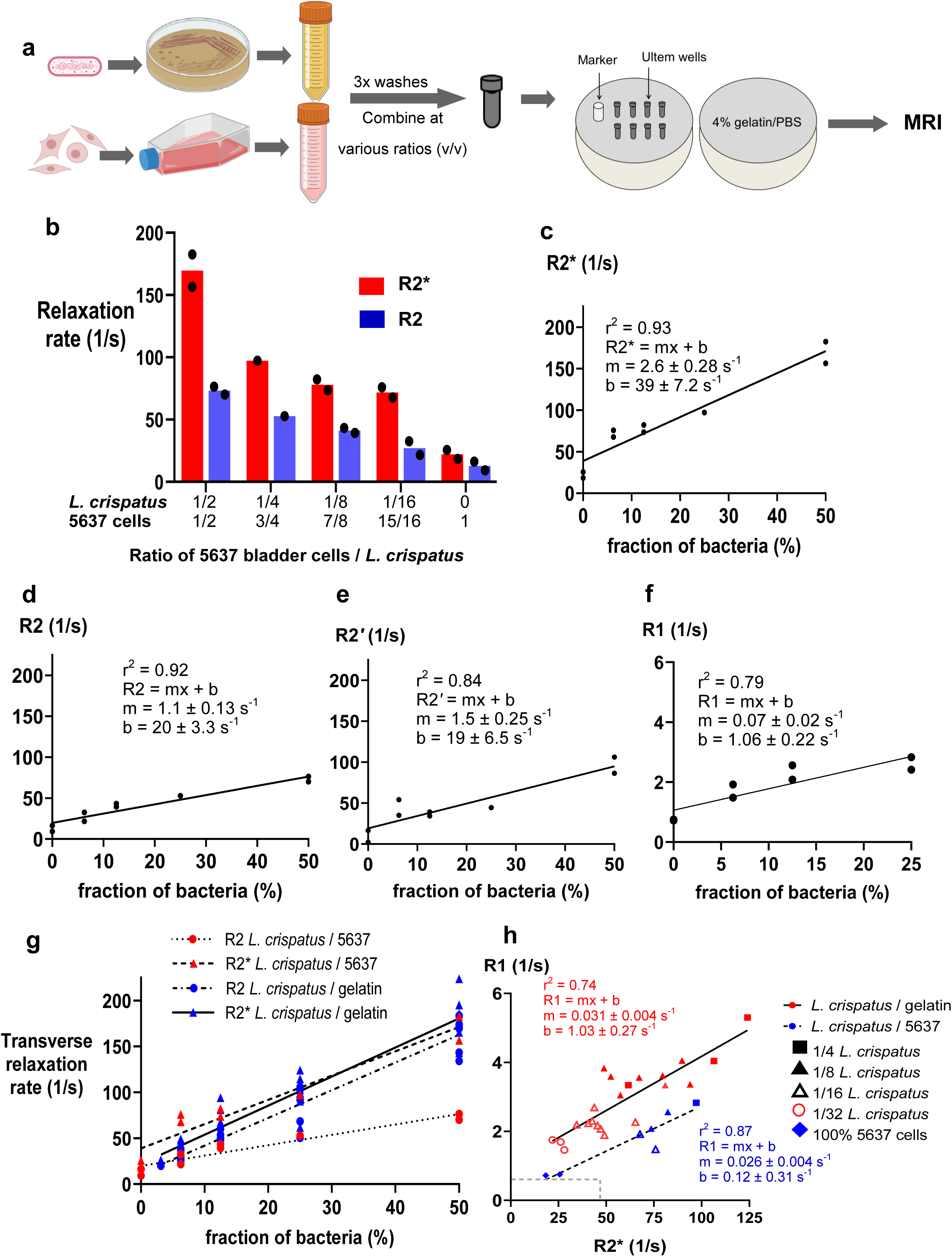
Human bladder cells attenuate R2 transverse relaxation rates of *Lactobacillus crispatus* ATCC33820. **a)** *Lactobacillus crispatus* and human 5637 bladder cells were cultured and washed separately before mixing to serially dilute *L. crispatus* with increasing numbers of bladder cells. The resulting mixtures were then loaded into wells and mounted in a spherical cell phantom for MRI at 3T. **b)** Bar graphs show the influence of increasing proportions of bladder cells on *L. crispatus* transverse relaxation rates. Irrespective of dilution, R2 (blue bars) comprises approximately half of the R2* signal (red bars). **Cc-f)** Based on Pearson’s correlation analyses, fraction of *L. crispatus* is strongly positively correlated to R2* (c), R2 (d), R2′ (e), and R1 (f) measurements. **g)** The graph of cumulative data compares R2 (circles) and R2* (triangles) against the fraction of bacteria. Blue symbols denote *L. crispatus* diluted in gelatin alone; red symbols denote mixed samples of *L. crispatus* and bladder cells. Lines demonstrate the linear regression within each sample type and MR measure. **h)** The graph of cumulative data compares paired R1 and R2* measurements of *L*. *crispatus* dilutions. Red symbols show *L. crispatus* diluted in gelatin, with a solid black linear regression curve described by the equation in red. Blue symbols show *L. crispatus* reduced in number by the addition of 5637 bladder cells, with a dotted black linear regression curve described by the equation in blue. Dilution factor is denoted by symbol shape: 1/4, square; 1/8, closed triangle; 1/16, open triangle; 1/32, open circle. Values for bladder cells alone are indicated by a blue diamond. Broken gray lines show the average R1 (0.67 s^-1^) and R2* (39.3 s^-1^) in the healthy human bladder.

The R2* values from two samples of human 5637 bladder cells (Fig. 5b; 18.4 and 25.6 s^- 1^, right-most red bar) are in the range of many other mammalian cell types. For comparison, in human melanoma MDA-MB-435 cells the measured value for R2* is 13.70 ± 3.07 s^-1 43^ and in multi-potent mouse embryonic adenocarcinoma P19 cells R2* is 13.69 ± 0.59 s^-1 42^. These are nevertheless much lower than transverse relaxation rates of lactobacilli, with mean R2* of 210 ± 20.8 s^-1^ when diluted 1/2 (Fig. 4b). Whether in gelatin (Fig. 4, f-h) or reduced in concentration by the addition of different amounts of bladder cells (Fig. 5, c-f), fraction of *L. crispatus* cells is strongly and positively correlated to transverse and longitudinal relaxation rates. Comparing these linear regression slopes (Fig. 4, f and g vs Fig. 5, c and d), demonstrates that R2 but not R2* is decreased in the presence of bladder cells (Fig. 5g; r^2^ > 0.82 for all Pearson correlation coefficients; refer to Fig. 4, f and g). The linear regression of R2 vs. *f* for bacteria / gelatin is significantly different than that of bacteria / bladder cell mixtures (p < 0.001; Sum-of-squares F-test, F(6,68)=17.98). Accordingly, the ratios of R2/R2* are significantly lower for *L. crispatus* in the presence of bladder cells than for those bacteria diluted in gelatin alone (Table 2; p < 0.001; Mann-Whitney *U*-test, W=437 P < 0.0001).

**Table 2.**
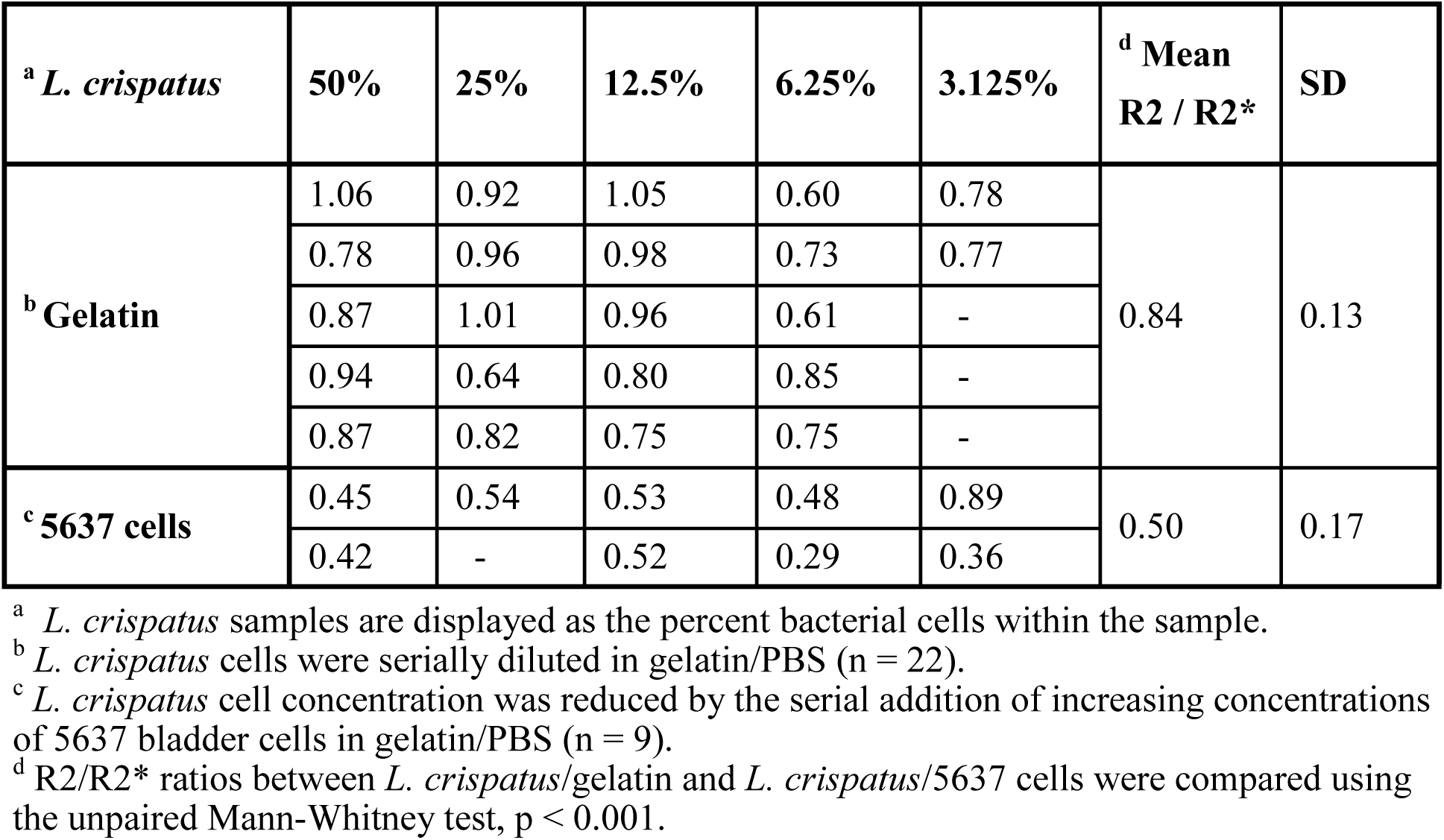
R2/R2* ratios for L. crispatus ATCC33820 dilutions

Longitudinal relaxation rates for mixtures of *L. crispatus* and bladder cells were also linearly correlated with fraction of bacteria (Fig. 5f, Supplementary Table 4). A scatter plot of R1 vs R2* demonstrated that both measures are strongly correlated; increase linearly in the presence and absence of bladder cells; and even at 1/32 dilutions of *L. crispatus*, the signal is well above tissue background as assessed in male and female bladders (Fig. 5h). The same was true for R1 vs. R2 plots (Supplementary Fig. 4). As expected, relaxation rates in human bladder cell phantoms (blue diamonds, Fig. 5h) are within the range measured for bladder tissue (grey dashed line) and validate the cell phantom model.

## Discussion

The healthy lower urinary system has comparatively low microbial diversity yet appreciable bacterial density (∼ 100 million CFUs/mL in the vagina): features that enable imaging studies in this area. We thus assessed MR relaxation rates of species that have been detected at this site, using MRI to explore the bacterial detection limit. Examination of 13 distinct bacterial isolates revealed that many species have unique MR signatures, reflecting in part their intrinsic capacity for regulating Fe and Mn content. By examining bacteria with extraordinarily high transverse and longitudinal relaxation rates, MR measures were related to the quantity of live bacterial cells and hence to an estimate of CFUs needed to non-invasively detect bacterial infection by MRI.

Data obtained in a gelatin cell phantom indicate the feasibility of *in vivo* detection of select bacteria with MRI. Using 1/8 dilutions of *L. crispatus* for a conservative estimate, both transverse (R2* = 71.3 ± 6.10 s^-1^; R2 = 56.6 ± 3.50 s^-1^) and longitudinal (3.54 ± 0.10 s^-1^) relaxation rates should be useful for tracking less than 290 million cells per mm^3^ (11 billion cells / 38 mm^3^ MR slice). This sensitivity of detection using standard MR sequences at 3T is expected to improve with the application of ultrashort TE (UTE) ^45^, designed to capture the type of rapid decay of relaxation evident in species like *L. crispatus*.

Translation to future clinical applications involving MR detection of bacterial cells in the human host should be expected to depend on the possible additive or attenuating effects of multiple cell types and the potential for differentiating bacterial MR signatures within these complex environments. For *in vivo* regions of interest, the MRI voxel will generally contain both bacterial and mammalian cells since few bacteria within the human microbiota are free floating. Rather, bacteria commonly adhere to mammalian tissues via pili ^46^ or biofilm, the latter of which also form on abiotic materials such as catheters and stents ^47^. In addition, microbial diversity varies between each micro-environment ^48^, such that all clinical microbiota samples represent mixtures of various species living symbiotically.

In this work, we measured relaxation rates from representative two-component mixtures with the components being *L. crispatus* / gelatin in one mixture and *L. crispatus* / bladder cells in the other. Such systems are often represented by two compartment models. The MR behaviour of these systems depends on the fraction of water in each compartment as well as on the exchange rate of water molecules between compartments. Typically, the exchange rate is categorized into three regimes: fast, intermediate, and slow. Fast exchange refers to the case where the typical lifetime (or residence time) of water molecules in each compartment is short compared to the relaxation time being considered (T1 or T2; *i.e.*, 1/R1 or 1/R2).

The relaxation behaviour observed herein is consistent with fast exchange in two regards. (1) Signal decays are described by a single exponential function (rather than a two-component exponential), with the caveat that a separate exponential from *L. crispatus* could be too fast to be measurable here. (2) There is an approximately linear dependence of relaxation rate on the fraction of water in one compartment (Fig. 5g). However, extrapolating the regression lines of these two systems to estimate R1 and R2 values for *L. crispatus* alone (*i.e.*, *f* = 100%) provides very different relaxation rates (e.g., extrapolated R2 = 359 ± 27 s^-1^ for the gelatin / *L. crispatus* mixture and 130 ± 17 s^-1^ for the bladder cell / *L. crispatus* mixture). This suggests that the simple fast exchange model does not likely apply to both systems. We expect that the latter R2 value (130 ± 17 s^-1^) is an underestimate since it was within our measurable range. Also, we speculate that the exchange between compartments may be slower in the mixture with bladder cells, compared to the mixture with gel, due to the presence of mammalian cell membranes ^49^. The slower exchange would be associated with slower fluctuations of the microscopic magnetic field felt by water molecules moving through these spatial variations. This could potentially lead to greater signal reversibility by a refocusing pulse and thus lower R2 compared to R2* as we observed for this mixture with bladder cells.

While the two-compartment model is useful for initial consideration, it does not account for potential interaction of compartments. For example, *L. crispatus* contain high levels of paramagnetic ions (Mn) and the strong microscopic magnetic field variations produced would influence transverse relaxation (R2*, R2) of water within gel and bladder cells (*i.e.*, Mn in one compartment would affect transverse relaxation in the other compartment).

The high transverse relaxation rates of *L. crispatus* ATCC33820 may facilitate detection of this bacterium even at MRI field strengths lower than 3T, since R2 and R2* are generally proportional to field strength up to 3T ^50, 51^. Furthermore, given the moderate positive correlation between MR parameters and CFUs, *in vivo* imaging of *L. crispatus* and estimation of the number of live cells within an ROI should be feasible. For *L. crispatus*, fewer than 100 million CFUs / mm^3^ provides measurable MR relaxation rates that are greater than many mammalian tissues ^52, 53^.

Using the MR sequences reported herein, it would be nearly impossible to directly measure transverse relaxation of undiluted *L. crispatus* as its estimated R2 (359 ± 35.5 s^-1^) corresponds to a signal decay time (T2) of approximately 3 ms. While such a problem might rarely be encountered *in vivo*, in healthy individuals where bacteria are more homogeneously distributed within their niche(s) ^54^, future advancements in MR imaging of bacteria may benefit from sequences incorporating UTE or zero echo time (ZTE) to visualize the rapid signal decay of bacteria with large R2 ^45^. For example, human cortical bone has recently been imaged using both UTE and ZTE with TE as low as 8 µs to improve bone and microstructure imaging that is normally difficult to evaluate with many common MR sequences ^55^.

Microscopic magnetic field inhomogeneities represented by R2′ (the difference between R2* and R2) may arise from potential differences in the form of metals within a cell (*i.e.* protein-bound, sequestered, redox active) and may inform the relation between components of transverse relaxation. Total cellular iron content varied among bacteria; however, in all lactobacilli examined in this study elemental iron fell below the detection limit. This strategy by *Lactobacillus* spp. of replacing iron ^34, 39, 56^ with another metal ion cofactor like manganese may provide a growth advantage in the low iron environments of the urogenital tract. Interestingly, during menses when iron levels increase, some lactobacilli such as *Lactobacillus iners* continue to thrive while *L. crispatus* levels generally decrease and recover only as iron levels decrease ^57, 58^. Inter-species differences may relate to how efficiently the cell disposes of large amounts of iron that can lead to increased production of harmful free radicals which promote DNA damage and lipid peroxidation ^59^. On the other hand, low molecular weight molecules containing manganese play a role in blocking the production of reactive oxygen species and prevent lipid peroxidation, allowing lactobacilli to safely accumulate high levels of intracellular Mn ^39, 60^. Consistent with this, cultured lactobacilli thrive in growth medium containing approximately 1000-fold more manganese than standard lysogeny broth (LB) ^39, 61^. The possibility that manganese is also externally bound has not been previously reported but cannot be ruled out. While manganese is a required cofactor ^36, 56^, high Mn in our samples could reflect metal externally bound to the cell membrane or peptidoglycan layer as well as internal stores. Lactobacilli are known to bind heavy metals like cadmium, lead and mercury ^26^. However, within lactobacilli the presence of both conserved manganese transporters (*e.g.*, MtsA ^36, 62^) and manganese-dependent metabolism is consistent with a high intracellular content of elemental Mn.

High Mn content in lactobacilli may be promoting high MR relaxation rates ^63^; however, in the species examined, level of Mn does not appear to be the only influence on transverse relaxation. For example, *L. gasseri* ATCC33323, with R2 of 75 s^-1^, has 50-fold less Mn than *L. rhamnosus* GR-1, with R2 of 48 s^-1^ (1 µg Mn / mg protein vs 53 µg Mn / mg protein, respectively). Moreover, total elemental manganese content of *L. rhamnosus* and *L. crispatus* are similar (53 vs 63 µg Mn / mg protein, respectively) despite a huge difference in R2 (48 vs an estimated 359 s^-1^, respectively). Since higher cellular manganese content does not necessarily equate with higher transverse relaxation, and elemental iron content is negligible, other factors are at play. These factors may include, but are not limited to, bacterial size, morphology, density of cells within a ROI, water content, and presence of additional MR sensitive elements, including nickel, cobalt and gadolinium. Although gadolinium is not typically found in bacteria, the genomes of some species like *Salmonella enterica*, *Campylobacter jejuni* and *Bacillus subtilis* encode nickel and/or cobalt transport proteins ^64^.

The notion that bacteria themselves may behave as contrast agents has not been widely explored. However, the modelling of select bacteria as micron sized nanoparticles may be useful for optimizing MR sequences, capitalizing on the fast and slow exchange of water to distinguish cellular and tissue structures ^65^, and discriminating mammalian from microbial signals and perhaps one bacterial species from another. Both the type and concentration of metal ion(s) in distinct bacteria may turn each species into a distinct MR nanoparticle, affording new opportunities in microbial cell tracking by tuning MR sequences for the detection of specific bacteria. Nanoparticles consisting of commensal species with high manganese content may yet serve as useful alternatives to gadolinium chelates that provide T1 contrast at the risk of kidney toxicity. When used in an ingestion model to examine gut permeability ^66^, such bacterial nanoparticles might offer new insights into bacterial translocation in addition to exploiting both R1 and transverse relaxation rates.

With the possibility of longitudinally tracking bacteria *in vivo* using MRI, and its hybrid modalities like PET/MRI, the feasibility of non-invasively imaging infections and microbial therapies moves closer to clinical implementation. As a non-ionizing imaging platform, MRI is not only suitable for a wide range of patient care, from young to old, but also provides superior soft tissue resolution with the option of multi-parametric imaging. These molecular imaging options provide timely alternatives to standard care and should expand clinical studies of the microbiota and infectious diseases.

## Acknowledgements

The authors thank technologists John Butler and Heather Biernaski for MR support and Dr. Kait Al and Wongsakorn Kiattiburut for valuable discussion on principal component analysis.

## Author contributions

S.C.D, J.P.B and D.E.G. conceived and designed the experiments. S.C.D. performed the experiments. S.C.D. and N.G. analysed the data. J.P.B., D.E.G., F.S.P. and R.T.T. contributed resources and funding to this study. D.E.G. and S.C.D. wrote the paper. All authors reviewed and edited the manuscript.

## Supplementary information

Supplementary data is attached.

## Declaration of interests

S.C.D, R.T.T., F.S.P., J.P.B. and D.E.G. are inventors on a patent relating to imaging of bacteria. Goldhawk, D.E., Burton, J.P., Silverman, M.S., *Donnelly, S.C., Thompson, R.T., Zhang, M. and Prato, F.S. Biomedical imaging of bacteria and bacteriophage. Multi-Magnetics Inc. (MMI) 2020 U.S. Provisional Patent Application # 63/016,68.

## Methods

### Reagents

Unless otherwise noted, all reagents were from Thermo Fisher Scientific, Mississauga, Canada and Sigma-Aldrich, Oakville, Canada.

### Bacterial culture

*Escherichia coli* strains were grown in LB for 16-20 h at 37°C with shaking. Staphylococci were grown in Brain-Heart Infusion (BHI) broth for 16 h at 37°C with shaking, or, when noted, in iron depleted RPMI-1640 medium for 24 h at 37°C with shaking. *Pseudomonas aeruginosa* PA01 was grown in LB for 16 h at 37°C with shaking. All lactobacilli were grown in deMan, Rogosa and Sharpe (MRS) broth anaerobically for 24 h at 37°C. *Proteus mirabilis* 296 and *K. pneumoniae* 280 were grown in LB, while *E. faecalis* ATCC33186 was grown in BHI, all anaerobically for 24 h at 37°C.

### Bacterial quantification

After bacteria were grown overnight, 200 µL of each culture was placed in a 96-well plate and measured in an Eon BioTek plate reader (Biotek, Winooski, USA) with Gen5 2.01 software to obtain OD_600_ measurements. This aliquot of each culture was then serially diluted 1/10 down to 10^-7^ and 10 µL volumes were plated in triplicate on LB/agar or BHI/agar. *Proteus mirabilis* samples were always plated on 6% non-swarming LB/agar. For lactobacilli, 5 µL of each dilution was plated in triplicate on pre-warmed MRS/agar plates. After drying, plates were inverted and incubated either aerobically or anaerobically (depending on the bacterium) at 37°C for 12–24 h before counting the number of CFUs. Equation 1 was used to calculate the total number of CFUs within the full culture based on the total culture volume and volume plated.

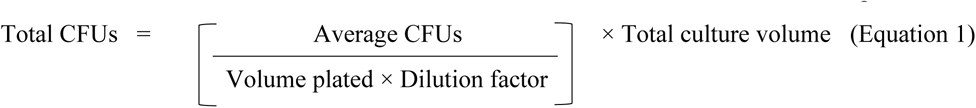

The remaining culture was routinely pelleted at 4500 *× g* for 10 min, washed three times with PBS, resuspended as a ∼75% cell slurry, and loaded into Ultem wells (Fig. 1a) by centrifugation at 4500 *× g* for 10 min. Supernatant was removed and more of the cell slurry was added and centrifuged until the Ultem well was full. Total number of CFUs within the wells were estimated using Equation 2 based on CFU counts.

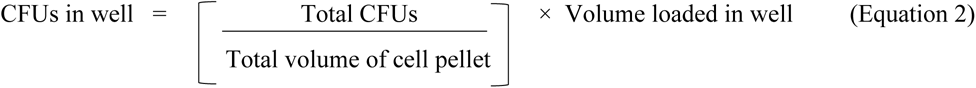

The number of CFUs within the MR slice (Fig. 1b) constitute ⅓ of the total CFUs loaded within the well (slice thickness is 3 mm and the total inner height of the Ultem well is 9 mm).

For *L. crispatus* dilutions, cells were cultured and quantified as described above but final cell pellets were diluted 1/2 in 4 % gelatin / PBS before serially diluting 1/2 down to a final dilution of 1/32. Diluted cells were mixed gently; loaded into Ultem wells; and immediately placed at 4°C to solidify before mounting in the gelatin phantom.

### Tissue culture

Human bladder epithelial cells (ATCC 5637) were cultured at 37°C and 5% CO_2_ in RPMI-1640 medium containing 10% FBS and 4 U/mL penicillin/4 μg/mL streptomycin. For passaging, plates were washed twice with phosphate buffered saline pH 7.4 (PBS) prior to agitating in 0.05% trypsin/ethylenediaminetetraacetic acid (EDTA) and incubating at 37°C for 10 min before triturating. Harvested cells were centrifuged at 550 *x g* for 10 min at 10°C, then washed with PBS and resuspended in fresh media for plating.

### Bladder cells and *L. crispatus*

Human ATCC 5637 bladder cells were cultured as described above in 12 x 150 mm plates; harvested; and combined into one sample for the serial dilutions described below. At harvest, cells were collected into one tube, centrifuged to remove medium, and washed three times with PBS before obtaining cell counts by hemacytometry. *Lactobacillus crispatus* ATCC33820, collected as described above, including plating for CFU determination. A suspension of *L. crispatus* was serially diluted ½ with a suspension of bladder cells using the following ratios of mammalian:bacterial cells (v/v): 15:1 (93.75%/6.25%), 7:1 (87.5%/12.5%), 3:1(75%/25%) and 1:1 (50%/50%). Samples of bladder cells alone were also prepared for MR analysis. Two biological replicates at each ratio were loaded into Ultem wells by centrifugation at 550 *× g*; removing supernatant and loading more sample until wells were full of the cellular mixture. Total number of bladder cells as well as bacterial cells were calculated as per Equation 2, factoring in the ratio of each cell type.

### Magnetic resonance imaging in a cell phantom

Cells were loaded into Ultem wells as described above prior to mounting in a 9 cm spherical MR phantom made of 4% gelatin (porcine type A)/PBS. MR phantoms were scanned at 3T on a Biograph mMR (Siemens AG, Erlangen, Germany), adapting previously developed sequences ^43^ to acquire longitudinal and transverse relaxation rates.

A single slice with slice thickness of 3 mm (Fig. 1b) and field of view (FOV) of 120 x 120 mm was used for all image acquisitions. An inversion recovery (IR) spin echo sequence was used to acquire R1 (R1 = 1/T1) measurements. Matrix size of 128 × 128 mm^2^ give a voxel size of 1.5 × 0.9 × 0.9 mm^3^. Repetition time (TR) was 4000 ms and inversion times (TI) were 22, 200, 500, 1000, 2000 and 3900 ms. The scan time for the R1 measurements was approximately 39 min.

A single echo spin echo (SE) sequence was used (Fig. 1c) was applied for R2 (R2 = 1/T2) measurements and a multi-echo gradient echo (GRE) sequence for R2* (R2* = 1/T2*) measurements. The matrix size for both sequences was 192 × 192 mm^2^ leading to a voxel size of 1.5 × 0.6 × 0.6 mm^3^. For SE, measurements were obtained at TE of 13, 20, 25, 30, 40, 60, 80, 100, 150 and 200 ms; TR – TE was held fixed at 2000 ms ^67^. For the GRE sequence, measurements were obtained at TE 6.12, 14.64, 23.16, 31.68, 40.2, 50, 60, 70 and 79.9 ms; TR was 2000 ms; number of averages = 4 and flip angle was 60°. The scan times for R2 and R2* measurements were approximately 61 min and 26 min respectively.

Longitudinal and transverse relaxation rates were determined using custom software developed in Matlab 7.9.0 (R2010b). For R2 and R2*, this software was used to select a 21- voxel ROI within the sample, avoiding the wall of the Ultem well. Signal intensity within the ROI was determined at each TE and decay curves were plotted using Prism software (GraphPad; San Diego, CA) version 9.3.0, and a single exponential line of best fit. R2′ was calculated by subtraction (R2* - R2 = R2′). For R1, mean signal intensities at each TI were taken from a 9-voxel ROI within the sample. Signal intensities at each TI were plotted using custom Matlab code developed in Matlab 2019 and the standard inversion recovery equation was applied for determination of R1. Longitudinal relaxation rates with uncertainty greater than 25% were excluded from further analyses. Relaxation rates were reported as the mean +/- SEM using Prism software.

### *In vivo* magnetic resonance imaging

Healthy volunteers were scanned at 3T on a Biograph mMR to acquire longitudinal and transverse relaxation rates. All images were acquired in the sagittal plane.

T1 mapping was performed using the variable flip angle method ^68^ with two flip angles (2°, 10°) and spatial resolution of 0.7 × 1 × 3.3 mm (interpolated to 0.7 × 0.7 × 2 mm). Flip angles for the T1 map reconstruction were obtained using the “B1 mapping using the slice selective pre-conditioning RF pulse” method ^69^. The scan time for T1 imaging was approximately 1 minute.

T2* mapping was performed with a 2D multi-gradient echo sequence with 6 TE values ranging from 2.46 ms to 14.75 ms and spatial resolution of 1.1 × 1.5 × 4 mm (interpolated to 1.1 × 1.1 × 4 mm). The scan time for T2* imaging was approximately 7 minutes.

### Protein quantification and mass spectrometry

Bacteria were cultured as above and centrifuged at 4500 *x g* for 10 min at 20°C, washing three times with at least 10 mL PBS. Cell pellets were then collected in RIPA buffer containing Complete Mini protease inhibitor cocktail (Roche Diagnostic Systems, Laval, Canada) and lysed through five cycles of freeze-thaw. Protein concentrations were measured using the BCA assay and BSA as the standard ^70^. Absorbance at 562 nm was determined using the Eon plate reader.

For elemental iron and manganese analysis, samples containing 1–3 mg/mL of protein were evaluated using ICP-MS (Biotron Analytical Services, Western University, London, Canada). Briefly, samples were digested with nitric acid and heat, then filtered prior to mass spectrometry. The data reported here reflect total cellular iron or manganese content normalized to total amount of bacterial protein.

### Statistics

Statistical analyses were performed using Prism software. All replicates shown reflect biological replicates. MR relaxation rates and metal content of bacteria were analyzed using one-way ANOVA, where significance was defined at α = 0.05, followed by Tukey’s test. Non-parametric data from ICP-MS was assessed using the Kruskal-Wallis test with uncorrected Dunn’s multiple comparisons. Differences between R2 and R2* of species were analyzed by a two-tailed paired t-test.

For principal component analyses, covariance in mean MR and mean ICP-MS measures for bacterial strains was examined using Prism software. Data were standardized and variables examined were R2, R1, [Fe] and [Mn]. Eigenvalues for all principal components shown are greater than 1.0 and each set of principal components explained at least 85% of variance in the data. Follow up principal component analyses were completed after removing statistical outliers (Fig. 3d and Supplementary Table 3). Correlation analyses were completed using a one-tailed Spearman’s correlation at α = 0.05 with Bonferroni correction for multiple comparisons.

Comparison of R2 vs. *f* and R2* vs. *f* linear regressions for bacteria diluted in gelatin or bladder cells was done using an extra sum-of-squares F-test. Ratios of R2/R2* were compared using the unpaired two-tailed Mann-Whitney *U*-test to compensate for large differences in sample size between groups.

## Data availability

The data upon which this study was based are available from the corresponding author upon request.

## Code availability

Some data analysis was performed using MATLAB (Mathworks). These codes are available from the corresponding author upon reasonable request.

---

**Supplementary Figure 1.**
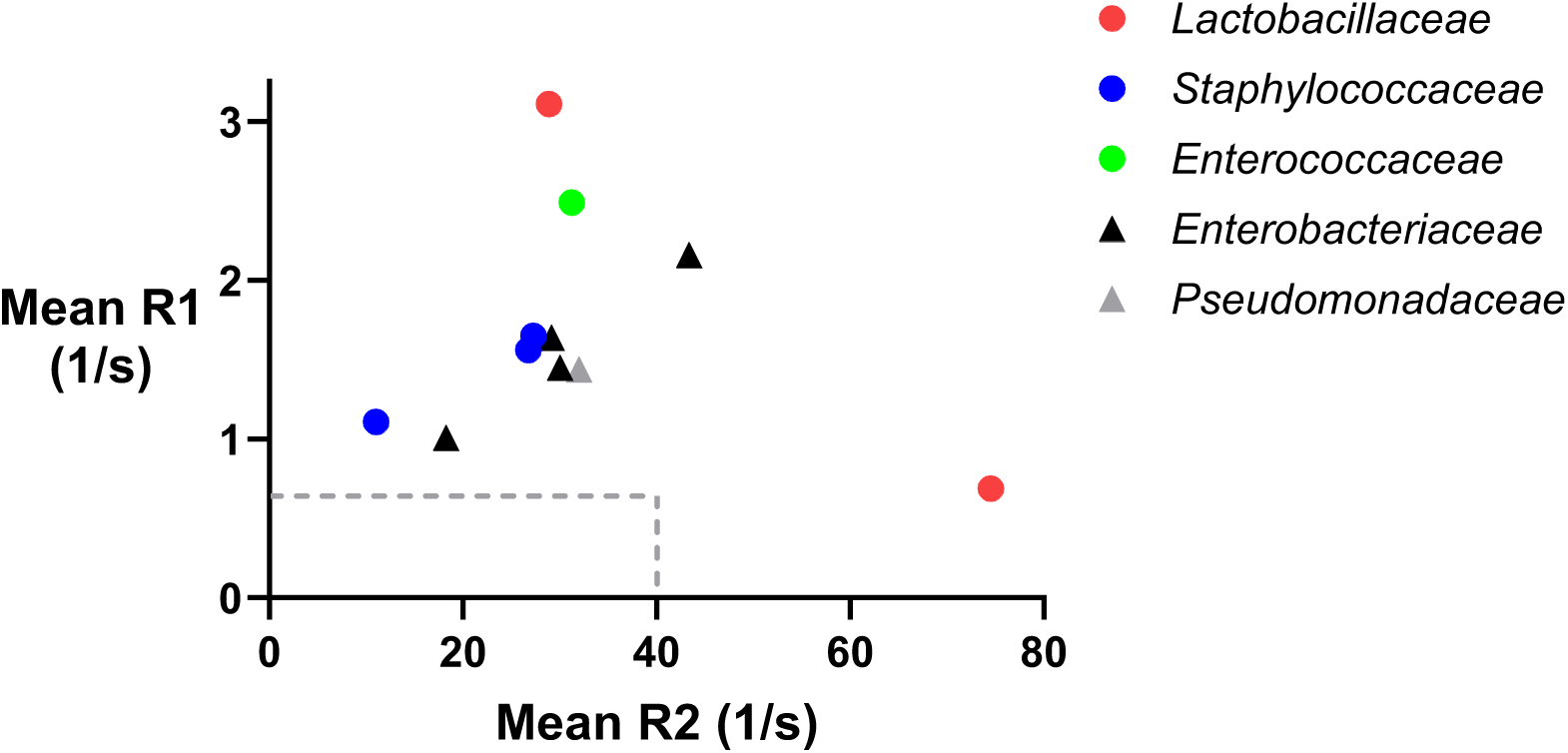
R1 vs. R2 of various bacterial species. Cultured bacteria were harvested, washed, and loaded into Ultem wells placed within a gelatin phantom and scanned at 3T by MRI. A scatter plot of R1 vs R2 shows the mean relaxation rate measurements of bacteria (n = 3–5). Gram positive species are denoted by circles and gram-negative species by triangles. Bacteria are grouped into their respective families as indicated by symbol colour: red, *Lactobacillaceae*; blue, *Staphylococcaceae*; green, *Enterococcaceae*; black, *Enterobacteriaceae*; gray, *Pseudomonadaceae*. Broken gray lines show average R1 (0.67 s^-1^) and R2* (39.3 s^-1^) in the healthy human bladder.

**Supplementary Figure 2.**
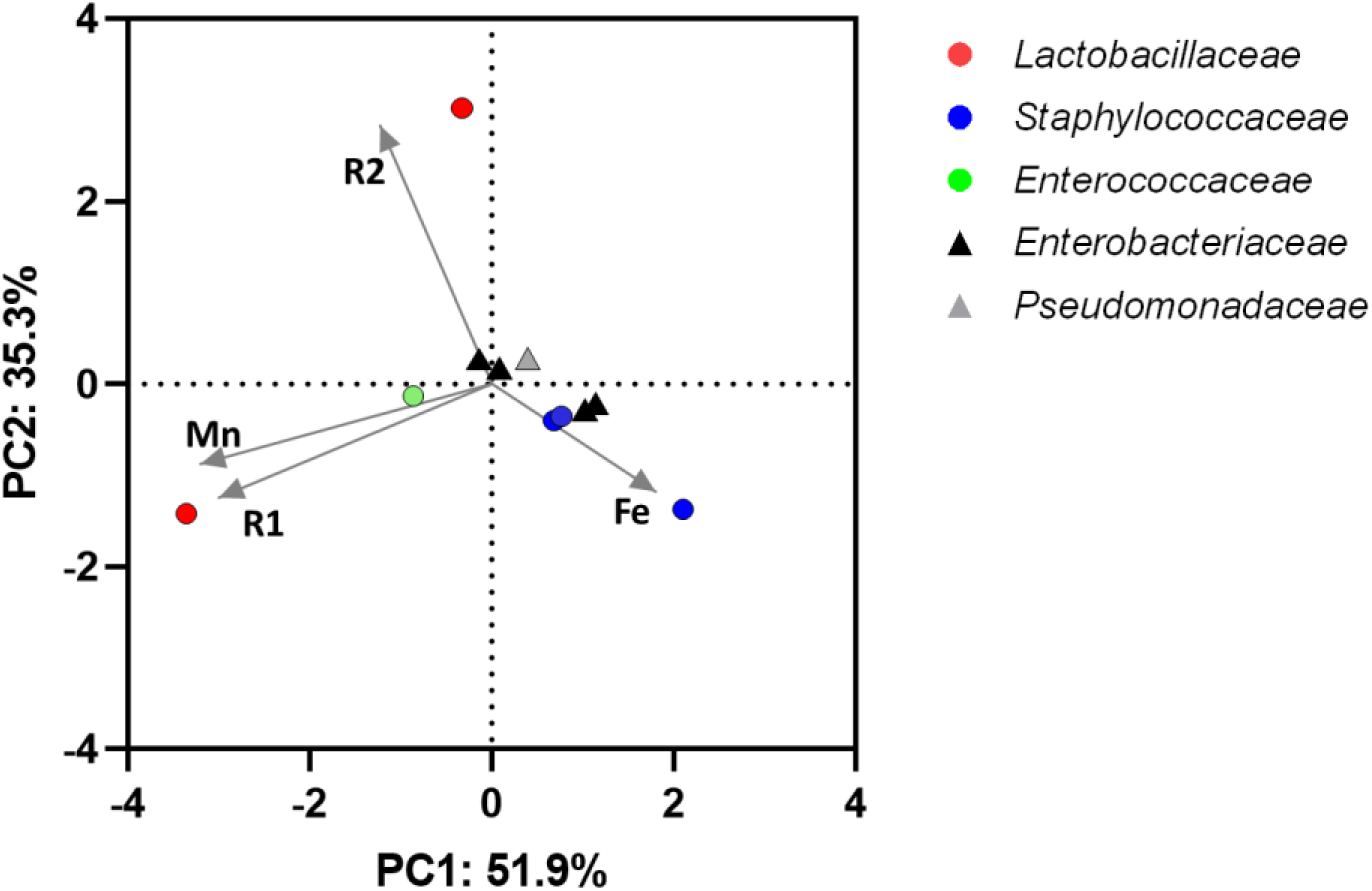
Principal component analysis of mean MR and ICP-MS measures in bacteria. The distance between samples on the plot represents differences in bacterial metal handling and MR measures, with 87.2% of total variance being explained by the first two components. The association of variables are depicted by the direction of the gray arrows. Each coloured point represents a separate bacterial strain or species as the mean of 3-5 replicates of each variable measurement. Points are coloured by bacterial family: red, *Lactobacillaceae*; blue, *Staphylococcaceae*; green, *Enterococcaceae*; black, *Enterobacteriaceae*; gray, *Pseudomonadaceae*. Gram positive species are denoted by circles and gram-negative by triangles.

**Supplementary Figure 3.**
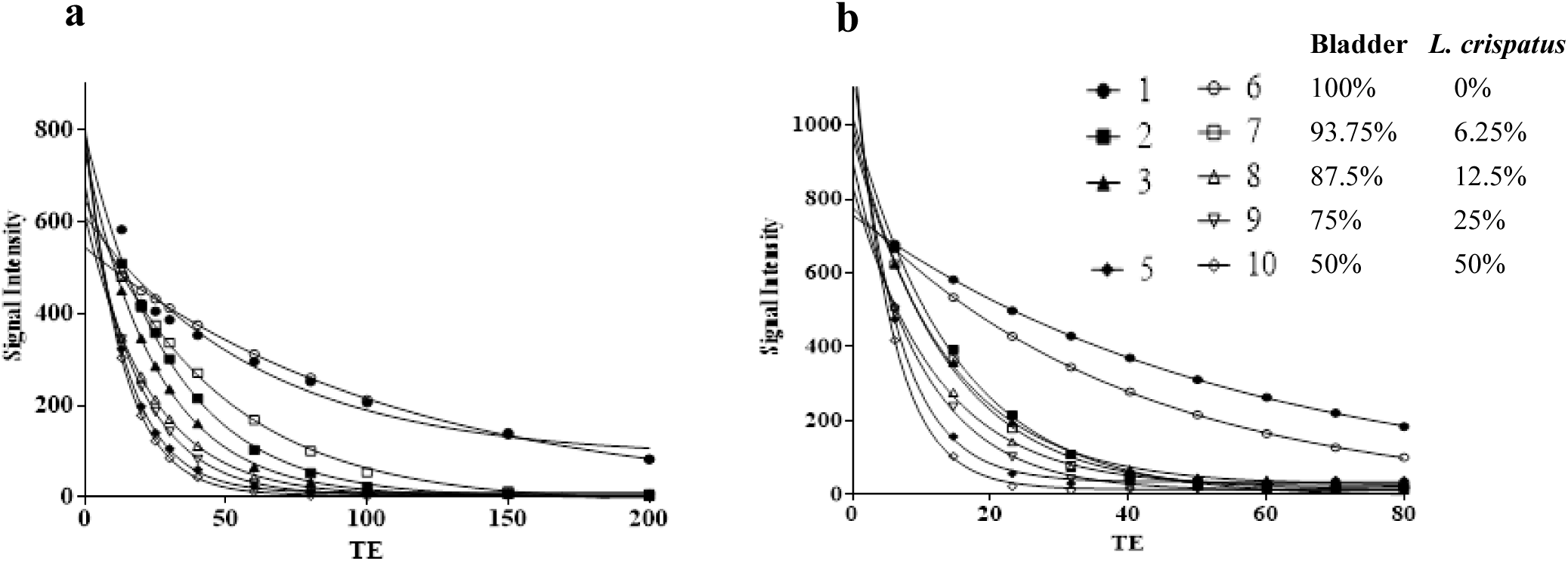
Signal intensity decays mono-exponentially with echo time (TE) in mixed samples of 5637 bladder cells and *L. crispatus* ATCC33820. The decay of T2 (a) and T2* (b) fits a mono-exponential curve, providing little or no distinction between mammalian and bacterial MR contributions. Open and closed circles (lines 1 and 6) show the signal decaying in bladder cells alone; open and closed squares (lines 2 and 7) show the signal decaying in bladder cells mixed with 1/16 *L. crispatus*; open and closed triangles (lines 3 and 8) show the signal decaying in bladder cells mixed with 1/8 *L. crispatus*; inverted triangles (line 9) show the signal decaying in bladder cells mixed with 1/4 *L. crispatus*; open and closed diamonds (lines 5 and 10) show the signal decaying with equal amounts of mammalian and bacterial cells (i.e. 1/2 *L. crispatus*).

**Supplementary Figure 4.**
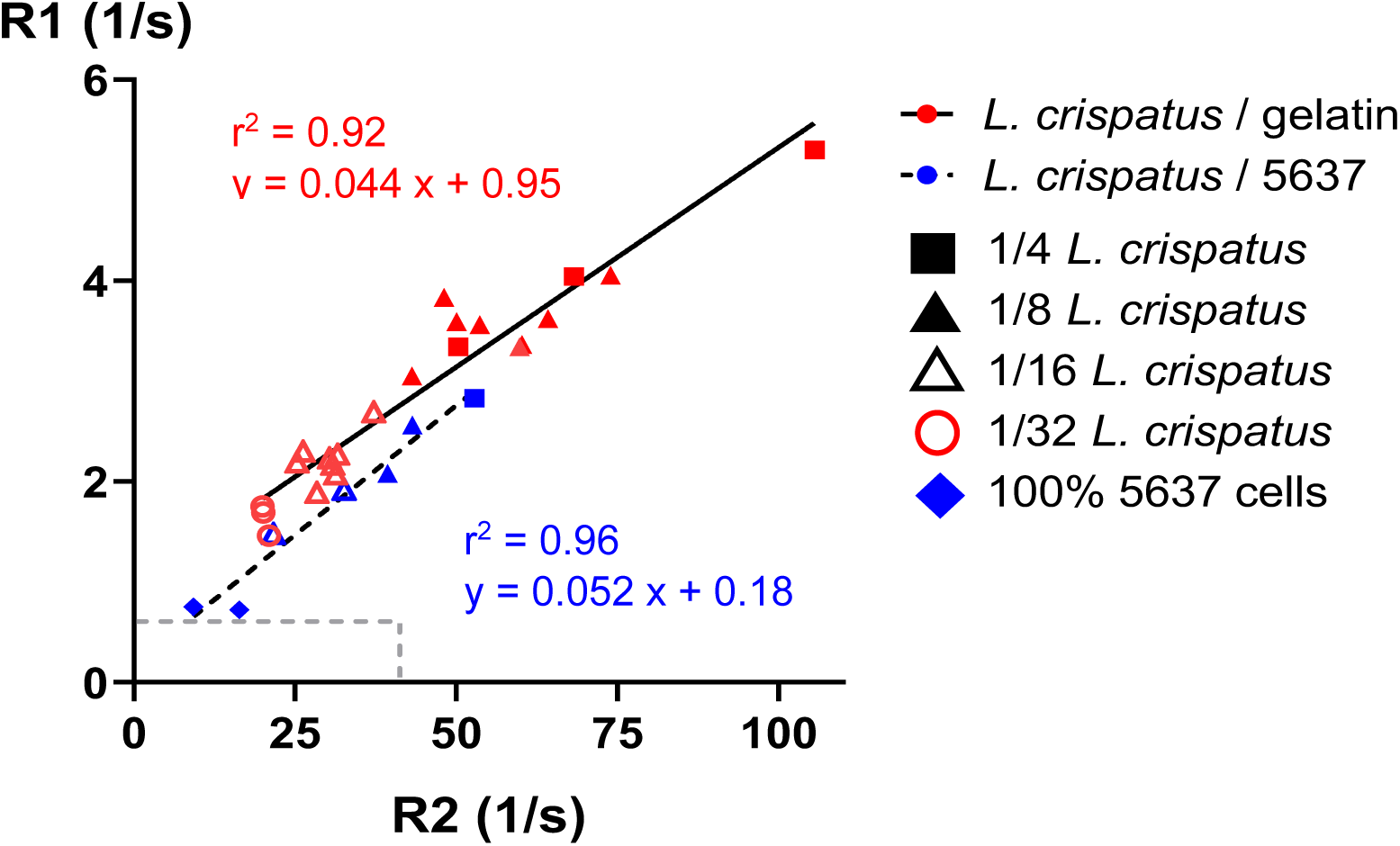
R1 vs. R2 of *L. crispatus* dilutions. *Lactobacillus crispatus* and human 5637 bladder cells were cultured and washed separately before mixing to serially dilute *L. crispatus* with either increasing numbers of bladder cells or in 4% gelatin/PBS. The resulting mixtures were then loaded into wells and mounted in a spherical cell phantom for MRI at 3T. The graph of cumulative data compares paired R1 and R2 measurements of *L*. *crispatus* dilutions. Red symbols show *L. crispatus* diluted in gelatin, with a solid black linear regression curve described by the equation in red. Blue symbols show *L. crispatus* reduced in number by the addition of 5637 bladder cells, with a dotted black linear regression curve described by the equation in blue. Dilution factor is denoted by symbol shape: 1/4, square; 1/8, closed triangle; 1/16, open triangle; 1/32, open circle. Values for bladder cells alone are indicated by a blue diamond. Broken gray lines show the average R1 (0.67 s^-1^) and R2* (39.3 s^-1^) in the healthy human bladder.

**Supplementary Table 1.** *In vivo* bladder imaging subjects and raw data

| Subject | Sex | Age | T1 (ms)<sup>a</sup> | SD (ms) | T2* (ms)<sup>b</sup> | SD (ms) |
| --- | --- | --- | --- | --- | --- | --- |
| 1 | M | 75 | 1852 | 214 | 23.9 | 6.2 |
| 2 | M | 59 | 1181 | 94.6 | 25.7 | 4.5 |
| 3 | F | 29 | 1489 | 121 | 22 | 8.8 |

<sup>a</sup> Mean T1 within these subjects was $1507 \pm 336$ ms, giving a mean R1 of $0.66 \pm 0.12$ s<sup>-1</sup>
<sup>b</sup> Mean T2\* within these subjects was $23.9 \pm 1.85$ ms, giving a mean R2\* of $41.8 \pm 3.55$ s<sup>-1</sup>

**Supplementary Table 2.** Spearman’s correlation for bacterial MR and ICP-MS measures of all urinary isolates examined ^a^

|  | **R2** | **R1** | **Fe** | **Mn** |
| --- | --- | --- | --- | --- |
| **R2** | 1.000 | 0.091 | -0.583 <sup>b</sup> | 0.091 |
| **R1** | 0.091 | 1.000 | -0.264 | 0.355 |
| **Fe** | -0.583 <sup>b</sup> | -0.264 | 1.000 | -0.415 |
| **Mn** | 0.091 | 0.355 | -0.415 | 1.000 |

<sup>a</sup> Covariance between MR relaxation rates and ICP-MS elemental analysis, where 1 is highly correlated and 0 is poorly correlated; negative correlations are indicated by a minus sign
<sup>b</sup> $p = 0.03$ was deemed not significant after applying a Bonferroni correction

**Supplementary Table 3.** Spearman’s correlation ^a^ for bacterial MR and ICP-MS measures for urinary isolates, excluding outliers ^b^

|  | **R2** | **R1** | **Fe** | **Mn** |
| --- | --- | --- | --- | --- |
| **R2** | 1.000 | 0.429 | -0.405 | 0.119 |
| **R1** | 0.429 | 1.000 | -0.357 | 0.881 <sup>c</sup> |
| **Fe** | -0.405 | -0.357 | 1.000 | -0.143 |
| **Mn** | 0.119 | 0.881 <sup>c</sup> | -0.143 | 1.000 |

<sup>a</sup> Covariance between MR relaxation rates and ICP-MS elemental analysis, where 1 is highly correlated and 0 is poorly correlated; negative correlations are indicated by a minus sign
<sup>b</sup> *Lactobacillus gasseri*, *L. reuteri*, and *S. aureus* Newman were considered statistical outliers and were excluded from the analysis
<sup>c</sup> Significant at $p < 0.05$ after Bonferroni correction

**Supplementary Table 4.**
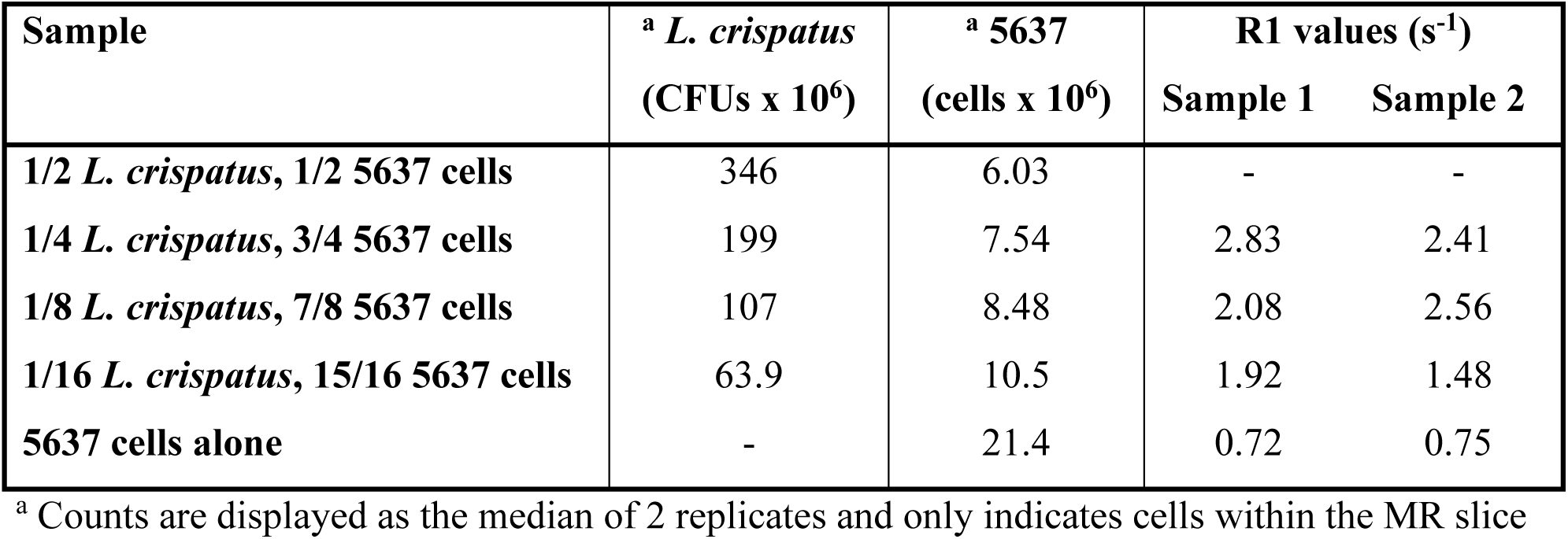
Longitudinal relaxation rates of mixed samples of 5637 bladder cell and *L. crispatus* ATCC33820

